# Zika virus infection preferentially counterbalances human peripheral monocyte and/or NK-cell activity

**DOI:** 10.1101/225102

**Authors:** Fok Moon Lum, David Lee, Tze-Kwang Chua, Jeslin J.L. Tan, Cheryl Y.P. Lee, Xuan Liu, Yongxiang Fang, Bernett Lee, Wearn-Xin Yee, Natasha Y. Rickett, Po-Ying Chia, Vanessa Lim, Yee-Sin Leo, David A. Matthews, Julian A. Hiscox, Lisa F.P. Ng

## Abstract

Zika virus (ZIKV) has re-emerged in the population and caused unprecedented global outbreaks. Here, the transcriptomic consequences of ZIKV infection were studied systematically firstly in human peripheral blood CD14^+^ monocytes and monocyte-derived macrophages with high density RNA-sequencing. Analyses of the ZIKV genome revealed that the virus underwent genetic diversification and differential mRNA abundance was found in host cells during infection. Notably, there was a significant change in the cellular response with crosstalk between monocytes and natural killer (NK) cells as one of the highly identified pathway. Immune-phenotyping of peripheral blood from ZIKV-infected patients further confirmed the activation of NK cells during acute infection. ZIKV infection in peripheral blood cells isolated from healthy donors led to the induction of IFNγ and CD107a — two key markers of NK-cell function. Depletion of CD14^+^ monocytes from peripheral blood resulted in a reduction of these markers and reduced priming of NK cells during infection. This was complemented by the immunoproteomic changes observed. Mechanistically, ZIKV infection preferentially counterbalances monocyte and/or NK-cell activity, with implications for targeted cytokine immunotherapies.

## Introduction

Zika virus (ZIKV) gained global attention in 2015-2016 when the virus suddenly re-emerged in the human population and caused major viral outbreaks across the world with a large disease burden (1). Although ZIKV has been causing sporadic outbreaks since it was first reported in Uganda >60 years ago (2), very little is known about the biology of the virus and the host response to infection. ZIKV is an arthropod-borne flavivirus that causes Zika fever — a disease that for the majority of patients has little or no symptoms (3). However, in severe cases, ZIKV infection may be responsible for neurological complications such as Guillain Barré Syndrome (GBS) in adults (4) and congenital fetal growth-associated anomalies in newborns (5). The host response to ZIKV infection may be one of the main drivers of the different disease phenotypes.

Recent studies have established that ZIKV can infect peripheral blood monocytes (6-9). However, despite ongoing intensive investigative efforts to understand ZIKV-related neuropathogenesis, knowledge regarding the mechanisms of ZIKV infection in peripheral immune cells is lacking. Given that ZIKV is transmitted into the dermis via the bite from a virus-infected mosquito, monocytes would be one of the first immune cells in the blood to interact with the virus when it reaches the circulatory system. Therefore, the interplay between ZIKV and monocytes will be crucial in determining the outcome of infection (10).

This study focused on characterising the primary *ex vivo* response of human donor blood monocytes and monocyte-derived macrophages (MDMs) to ZIKV infection. Systematically, RNA-sequencing (RNA-seq) was first used to identify and quantify the abundance of host messenger RNA (mRNA) and characterise viral RNA. This information was subsequently used to map the host response to ZIKV infection in the two different *ex vivo* cell types. These data also provided insights into the potential adaptation of the virus during viral replication in these cells. Immune-phenotyping of peripheral blood cells isolated from patients infected with ZIKV independently was executed to validate the predictions obtained from the differential gene expression analysis. Depletion of CD14^+^ monocytes in peripheral blood was then performed *ex vivo* to functionally understand the crosstalk between monocytes and priming of NK cells during ZIKV infection. Lastly, a multiplex assay was carried out to further understand host cell immunoproteomic changes during ZIKV infection. To our knowledge, this study is the first large-scale systematic investigation into the host cellular response to ZIKV infection in biologically relevant cells. This global analysis of the host immune response provides a novel understanding of the pathobiology of the virus, leading to the possibility of targeted therapeutic interventions in severe cases.

## Results

### ZIKV targets human peripheral blood monocytes and macrophages

CD14^+^ monocytes have been reported to be the main targets of ZIKV during infection (6-9). In this study, human primary CD14^+^ monocytes were first isolated from fresh peripheral blood mononuclear cells (PBMCs) to enrich this cell type to >90% of the total cell population (Figure 1A). In addition, isolated monocytes from the same donors were differentiated into monocytes-derived macrophages (MDMs) over 5 days (Figure 1B). Purified cells were then infected *ex vivo* with ZIKV and their permissiveness to ZIKV infection and growth was determined at 24 and 72 hours post-infection (hpi) (Figure 1A). The 24 hpi time point was chosen to represent the acute infection phase and the 72 hpi time point a stage by which a substantial host-virus interaction would have taken place (11). Data obtained showed that ZIKV infection of MDMs was more significant than infection of monocytes in all five donors (~40% compared to ~20% at 72 hpi, respectively) (Figure 1C). A decrease in viral load was observed in the virus-infected MDMs between the two time points, whereas the viral load remained consistent in infected monocytes over time (Figure 1D).

**Figure 1 :**
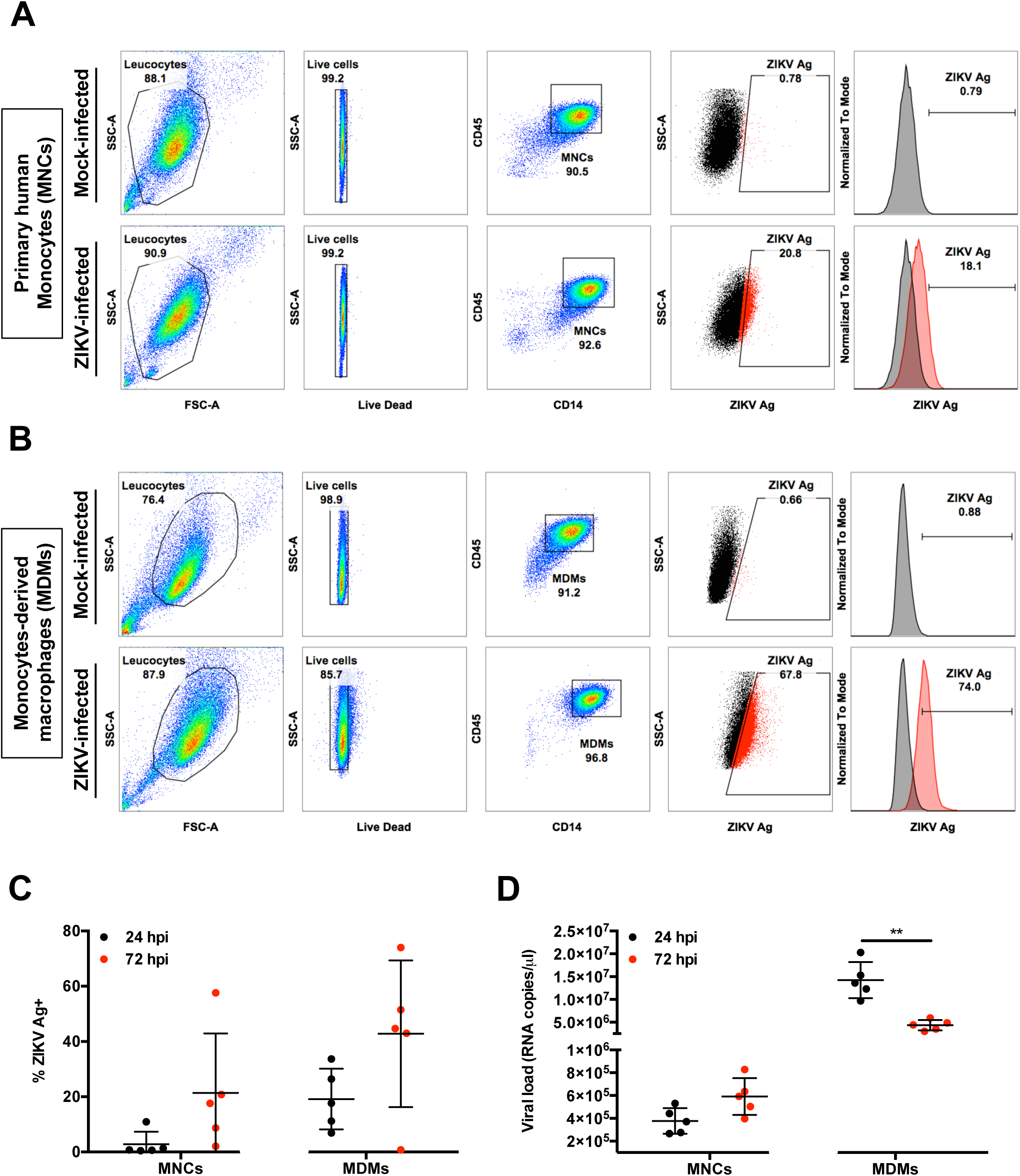
Primary human MNCs and MDMs are targets of ZIKV infection. Isolated human primary MNCs and MDMs (2×10^6^ cells each) were infected with ZIKV at MOI 10 and harvested at 24 and 72 hpi. Flow cytometry gating on **(A)** monocytes (MNCs) and **(B)** MDMs. Gating for positive infection was set using the mock-infected samples. For the dot plots, cells positive for ZIKV Ag are shown in red. For the histogram, ZIKV-infected samples (red) were overlaid on mock-infected samples (black). Compiled results for **(C)** infection (ZIKV Ag) and **(D)** viral load detected in MNCs and MDMs obtained from five healthy donors. All data are presented as mean ± SD. *\*P <* 0.05, by Mann Whitney *U* test, two tailed. Viral load data was not statistically significant between 24 and 72 hpi in MNCs by Mann Whitney *U* test, two tailed. Abbreviations: hpi, hours post-infection; MDM, monocyte-derived macrophage; MNC, monocyte; ZIKV, Zika virus; Ag, antigen.

In parallel, virus infection in PBMCs obtained from four healthy human volunteers also showed that CD14^+^ monocytes were the main immune subsets infected (Supplemental Figure 1).

### Genome variation in ZIKV during infection of the peripheral blood

In order to compare the amount of virus between the different cell types and determine whether ZIKV underwent genetic diversification during infection,viral sequence reads were mapped and compared to that of the progenitor virus stock (PF/ZIKV/HPF/2013). These data indicated that for MDMs, 4.53% and 0.43% of total sequence reads mapped to the ZIKV genome at 24 hpi and 72 hpi, respectively. While 24% and 0.8% of sequence reads generated from monocytes mapped to the ZIKV genome at 24 hpi and 72 hpi respectively. These observations are consistent with ZIKV viral load analysis, where higher levels of viral RNA were detected in MDMs (Figure 1).

Due to the inherent error-prone nature of viral RNA replication, nucleotide variants may become established in the viral genome during ZIKV infection in different cell types. To investigate this hypothesis, consensus genome information for each sample and the frequency of minor variants at each nucleotide position in the progenitor stock was determined and compared to the genome of virus present in the infected samples utilizing previously developed workflows (12,13). The ZIKV consensus genome sequence derived from the progenitor stock was 10,570 nucleotides in length and contained minor variants (as a measure of quasi-species) spread throughout the genome (Figure 2A). Of the 11 valid consensus sequences derived from the virus-infected samples, the virus recovered in cells from five donors (D1-D5) had the same consensus sequence as the input stock (PF/ZIKV/HPF/2013). However, some donor samples contained viral genomes that had additional nucleotide differences at six different positions (Table 1). These nucleotide differences (Table 1) were visualized as a maximum likelihood phylogenetic tree, where the input stock was used as the reference sample (Figure 2B). There were only eight high frequency transition mutations to choose from (log_10_8 = 0.9, see Figure 2A), increasing the likelihood of these changes appearing several times. Of these eight transition mutations, six appeared as major variants and thus changed the overall consensus sequence. The nucleotide positions of these six transition mutations (Table 2) indicated that all the changes in the consensus sequence were already present at relatively high frequency as minor variants in the input stock and were subsequently amplified during viral replication. Changes at nucleotide positions 2,815 and 4,211 were the most common, being found in ~35% reads mapping to the virus genome. Had these changes been found in ≥50% reads, they would have been classified as major variants and thus changed the consensus sequence (Table 2).

**Figure 2:**
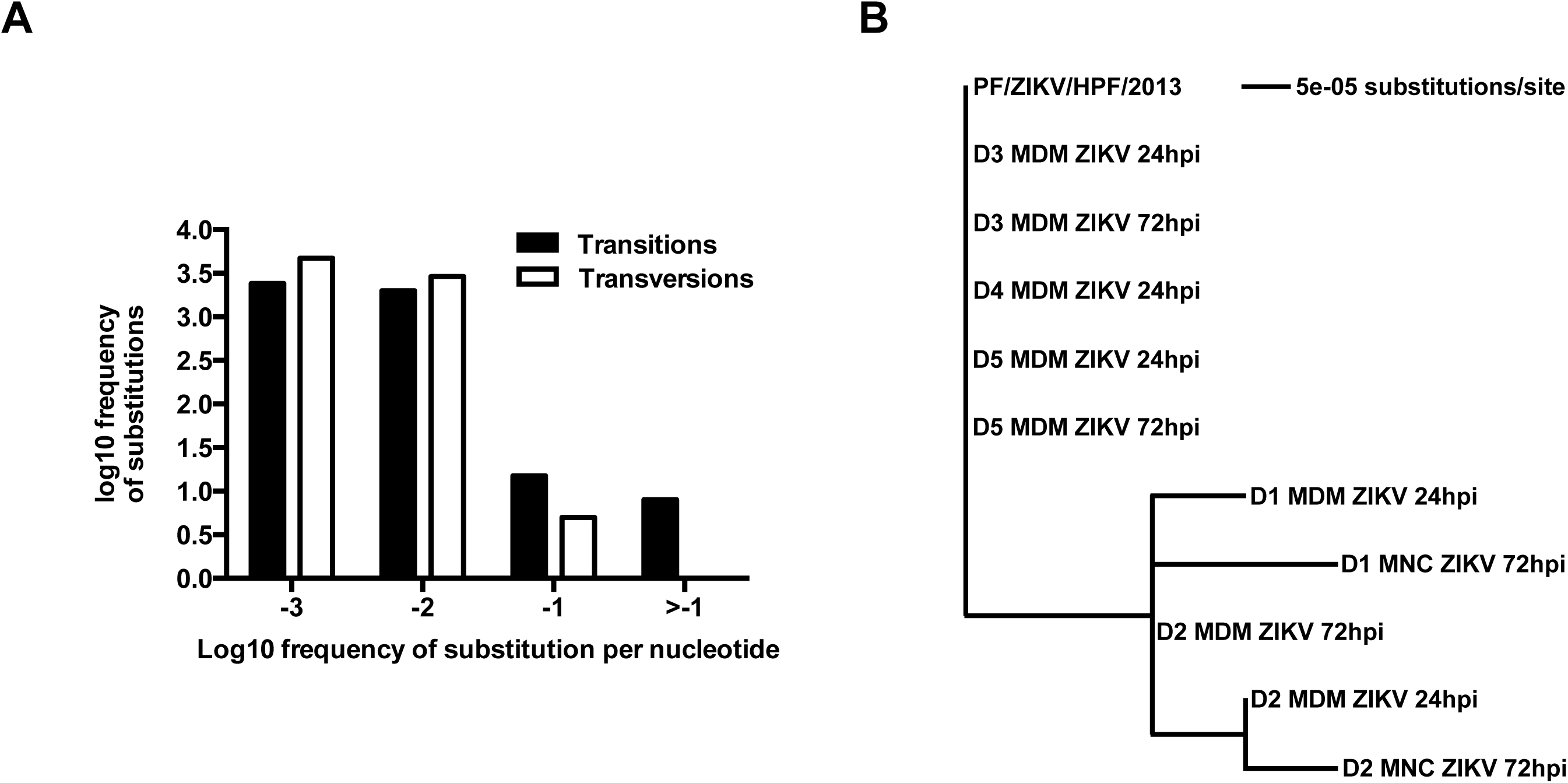
Phylogenetic analyses based on sample consensus sequences. **(A)** Frequency of ZIKV minor variants (transitions and transversions) recovered from infected human primary MNCs and MDMs isolated from five donors. Bin -3 is where ≤1/1000 reads show a specific change at an individual nucleotide position. Bin -2 is >1/1000 and ≤1/100 reads showing a difference. Bin -1 is >1/100 and ≤1/10 reads and Bin > -1 is >1/10 reads showing a change up to a logical limit of just under ½. **(B)** Phylogenetic tree generated from the alignment of consensus sequences of ZIKV RNA recovered from the same samples as described in **(A).** All samples included in the tree had a mean sequence coverage >10 at each nucleotide position. PF/ZIKV/HPF/2013 is the virus strain used for infection and denoted as the reference sample in this analysis. Abbreviations: hpi, hours post-infection; MDM, monocyte-derived macrophage; MNC, monocyte; ZIKV, Zika virus.

**Table 1:** Summary of nucleotide differences at specific genome positions. The phylogenetic tree shown in Figure 3B revealed specific nucleotide differences at 6 different positions within the consensus sequence. All samples included had a mean coverage of greater than 10. PF/ZIKV/HPF/2013 represents the virus used for the infection and therefore, denoted as the reference sample in this analysis. Abbreviations: MNC, monocytes; MDM, monocyte-derived macrophage. ^A^Note that minor variant file numbering of positions starts at 0 rather than 1.

| Position in consensus sequence<sup>A</sup> | Nucleotide difference | Sample(s) |  |
| --- | --- | --- | --- |
| 1904 | g → a | D1 MNC ZIKV 72hpi |  |
| 2673 | t → c | D2 MDM ZIKV 24hpi | D2 MNC ZIKV 72hpi |
| 2815 | t | D1 MDM ZIKV 24hpi | D2 MDM ZIKV 24hpi |
|  |  | D2 MDM ZIKV 72hpi | D1 MNC ZIKV 72hpi |
|  |  | D2 MNC ZIKV 72hpi |  |
| 2815 | c | D3 MDM ZIKV 24hpi | D3 MDM ZIKV 72hpi |
|  |  | D4 MDM ZIKV 24hpi | D5 MDM ZIKV 24hpi |
|  |  | D5 MDM ZIKV 72hpi | PF/ZIKV/HPF/2013 |
| 4211 | a | D3 MDM ZIKV 24hpi | D3 MDM ZIKV 72hpi |
|  |  | D4 MDM ZIKV 24hpi | D5 MDM ZIKV 24hpi |
|  |  | D5 MDM ZIKV 72hpi | PF/ZIKV/HPF/2013 |
| 4211 | g | D1 MDM ZIKV 24hpi | D2 MDM ZIKV 24hpi |
|  |  | D2 MDM ZIKV 72hpi | D1 MNC ZIKV 72hpi |
|  |  | D2 MNC ZIKV 72hpi |  |
| 10253 | t → c | D1 MDM ZIKV 24hpi | D2 MNC ZIKV 72hpi |
| 10472 | t → c | D1 MNC ZIKV 72hpi |  |

**Table 2:** Selected nucleotide positions from the minor variants file of the inoculum. Table showing the frequency distribution of minor nucleotide variants at six positions in the consensus sequence. Major variant (i.e. the consensus nucleotide) at each position is indicated by the nucleotide with the highest frequency. ^A^Note that minor variant file numbering of positions starts at 0 rather than 1.

| Position in consensus sequence<sup>A</sup> | Frequency of minor nucleotide variants |  |  |  |  |
| --- | --- | --- | --- | --- | --- |
|  | A | C | G | T(U) | D<br>(A/G/T) |
| 1904 | 0.11324 | 0 | 0.88601 | 0.00074 | 0 |
| 2673 | 0.00681 | 0.11779 | 0.000619 | 0.87476 | 0 |
| 2815 | 0.00876 | 0.64463 | 0 | 0.34659 | 0 |
| 4211 | 0.65122 | 0 | 0.34815 | 0.000626 | 0 |
| 10253 | 0.000891 | 0.11229 | 0 | 0.8868 | 0 |
| 10472 | 0.00108 | 0.00543 | 0.00108 | 0.99239 | 0 |

### Transcriptomic profiling reveals key cellular responses to ZIKV infection

RNA-seq was used to identify and quantify global mRNA abundance in ZIKV-infected peripheral monocytes and MDMs at 24 and 72 hpi. mRNA purified from 27 samples showed no signs of degradation and had sufficient read depth for inclusion in the analyses (Supplemental Figure 2A). For monocytes, mock and ZIKV-infected cells at both 24 and 72 hpi exhibited minimal changes in host transcript abundance. For MDMs, the abundance of transcripts that mapped to 1,736 and 545 genes at 24 and 72 hpi respectively, were significantly different (FDR < 0.05) between the mock and ZIKV-infected samples (Supplemental Figure 2B).

Ingenuity Pathway Analysis (IPA) was used to interrogate and group the differentially expressed genes into functional pathways (Figure 3A). A total of 169 pathways were identified, of which 27 were common in ZIKV-infected MDMs at 24 and 72 hpi. A further 106 pathways were unique to samples at 24 hpi (Supplemental Table 1), and 36 pathways were unique to samples at 72 hpi (Supplemental Table 2). This analysis found that genes associated with the interferon response were significantly upregulated at both time-points. In addition, signalling pathways involved in the pathogenesis of multiple sclerosis, and key pathways involved in monocyte-derived dendritic cell (moDCs) and NK cell processes were also shared between the two time points (Figure 3A). Overall, the top three common pathways activated in MDMs were interferon signalling, multiple sclerosis pathogenic pathways and crosstalk pathways between moDCs and NK cells (Figure 3A). The specific genes with the most abundant transcripts within these three pathways were analyzed, and when compared to the mock-infected controls were all increased in abundance after ZIKV infection (Figure 3B).

**Figure 3:**
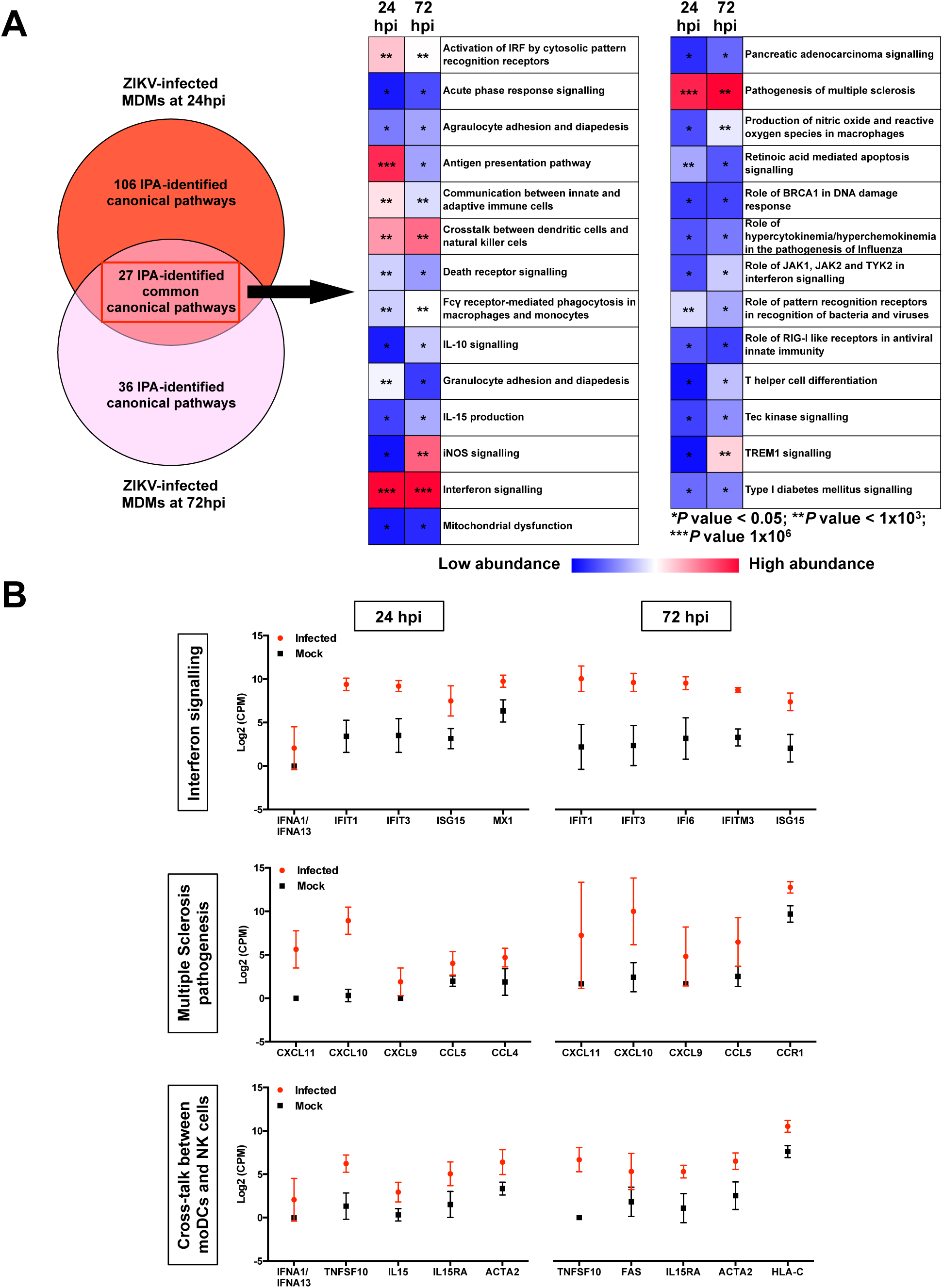
Transcriptomic profiling of host cells during ZIKV infection. Primary human MNCs and MDMs (2×10^6^ cells per infection) were infected with ZIKV at MOI 10 and harvested at 24 and 72 hpi for transcriptomic analysis by RNA-seq and then compared to mock-infected controls. **(A)** Venn-diagram illustrating the proportion of up-regulated signaling pathways identified by IPA in ZIKV-infected MDMs. Up-regulation intensity of the 27 common canonical pathways are shown in a heat-map. Stars within the boxes represent the calculated *P* values associated with each identified pathway, compared to the mock-infected samples. **(B)** The five most up-regulated genes within the top three signaling pathways at 24 and 72 hpi are shown: Interferon pathway, multiple sclerosis pathway and crosstalk between moDCs and NK cells. Data presented were obtained from a total of five donors. Abbreviations: hpi, hours postinfection; NK, natural killer; IPA, ingenuity pathway analysis; MDM, monocyte-derived macrophage; MNC, monocyte; moDCs, monocyte-derived dendritic cells; ZIKV, Zika virus.

### Virus-infected MDMs exhibit reduced cellular responsiveness

Transcriptomic profiles of various ZIKV-infected MDMs were compared to evaluate the transition of the cellular host response over the course of ZIKV infection. The percentage overlap of the identified transcripts between ZIKV-infected MDMs was assessed at 24 hpi and 72 hpi within the three targeted pathways described above (Figure 4). Interestingly, the percentage of overlapping transcripts identified at 72 hpi was lower for all three pathways, which may reveal a lower activation status of these pathways at this stage of the infection. The identification of different transcripts associated with 72 hpi may indicate the different signalling cascades present or activation status of these cells (Figure 4A). Global assessment of all identified transcripts revealed that transcripts mapping to 251 genes were in fact present in virus-infected MDMs at both time points. Transcripts that mapped to 1,485 genes were specific to 24 hpi, of which 54.81% exhibited greater abundance compared to the mock controls. By comparison, transcripts that mapped to 294 genes were unique to 72 hpi, with 63.36% of them having greater mRNA abundance compared to the mock controls (Figure 4B). Within the 251 common genes, transcripts mapping to 218 genes had a greater fold-change value compared to the mock-infected controls, indicating that these transcripts were increased in abundance in all ZIKV-infected MDMs. Further inquiry of these transcripts revealed that 60.1% of them were greater in abundance at 72 hpi compared to 24 hpi. Likewise, of the remaining transcripts that mapped to 33 genes and showed decreased abundance, 84.85% were further reduced at 72 hpi.

**Figure 4:**
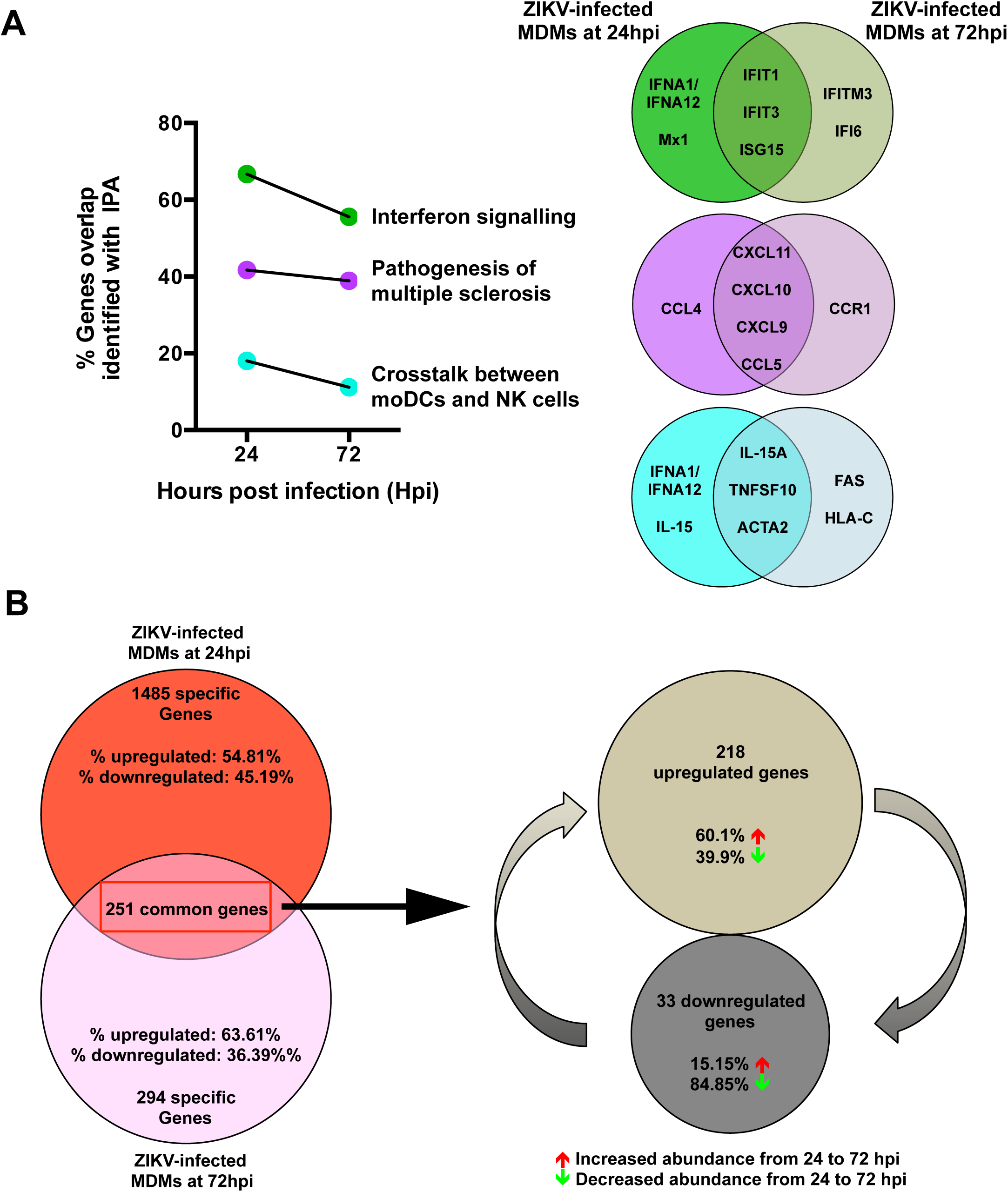
Transition of the host cellular response over the course of ZIKV infection. The host cellular response was analyzed and investigated by RNA-sequencing and significant transcriptomic differences were identified. **(A)** Transitional analysis (% genes overlapping) of the top three common canonical signaling pathways was determined using IPA of infected MDMs. Venn diagrams indicate the top five common and time-point specific genes associated with each canonical pathway. **(B)** Proportion of common and differentially expressed genes within ZIKV-infected MDMs at 24 and 72 hpi. Data presented were obtained from a total of five donors. Abbreviations: moDCs, monocyte-derived dendritic cells; hpi, hours post-infection; IPA, ingenuity pathway analysis; ZIKV, Zika virus; MDM, monocyte-derived macrophage.

### NK cells are activated in ZIKV-infected patients

IPA predicted robust crosstalk between NK cells and moDCs in peripheral blood upon *ex vivo* ZIKV infection (Figure 3-4). The IPA prediction that NK cells were activated in the peripheral blood of ZIKV-infected patients was, therefore, investigated by comprehensive immune-phenotyping of blood samples taken from ZIKV-infected patients. These patients were recruited from the first endemic ZIKV outbreak in Singapore in 2016 (7,14). Blood aliquots were obtained from ZIKV-infected patients (n=9) during the acute disease phase (between 1 and 7 days post-illness-onset), and were subjected to a whole blood staining protocol that targeted CD56^+^ cells, predominantly NK cells (15) (Figure 5A). Blood from healthy donors (n=5) was collected and processed in parallel as a control group. Gated cells were further grouped with the C-type lectin receptor CD94, giving three CD56^+^ populations: CD56^bright^CD94^hi^, CD56^dim^CD94^hi^ and CD56^dim^CD94^lo^ (16). The activation status of these populations was then assessed based on the percentage of each subset expressing CD16 and CD69 (Figure 5B). A higher level of CD16 was observed across all CD56^+^ subsets in ZIKV-infected patients compared to the healthy controls. A higher percentage of the subsets also expressed CD69 — a known cellular activation marker (17).

**Figure 5:**
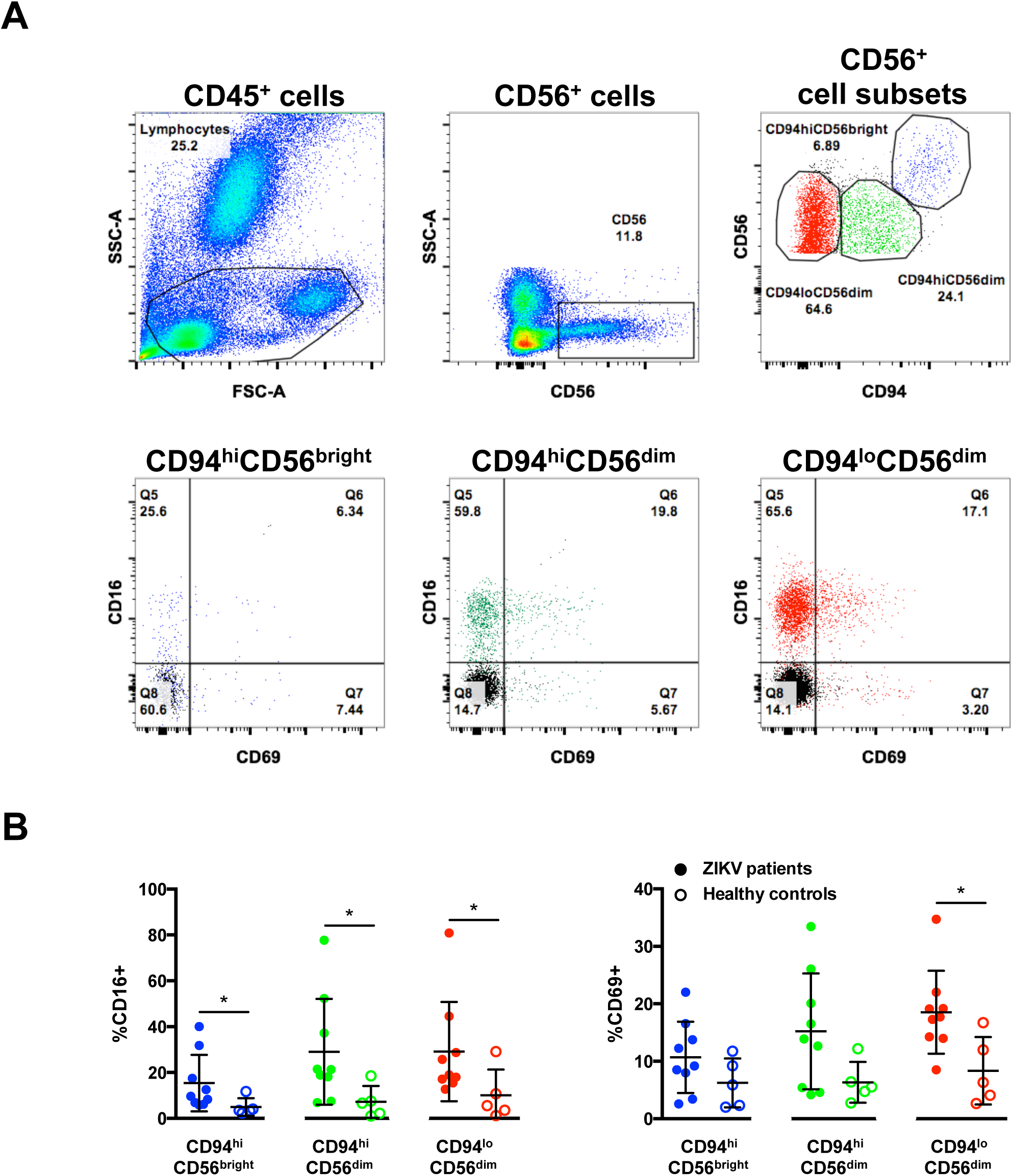
Activation of CD56^+^ cells in patients infected with ZIKV. (**A**) Gating strategy of CD56^+^ cell subsets and their expression of CD16 and CD69. CD56^+^ cells were first gated from CD45+ lymphocytes from peripheral blood mononuclear cells isolated from patients. These populations were further gated into three populations based on the expression of surface marker CD94: CD94hiCD56bright cells (blue), CD94hiCD56dim (green) and CD94loCD56dim (red). The data presented correspond to a representative patient infected with ZIKV. Cells from a healthy control are overlaid and depicted as the black population (Q8). (**B**) Compiled data on the percentage of gated subsets that are positive for CD16 (Q5 and Q6) and CD69 (Q6 and Q7). Patients (n=9) are depicted as filled circles, and healthy controls (n=5) are depicted as clear circles. All data are presented as mean ± SD. \**P* < 0.05, by Mann Whitney *U* test, two tailed. Abbreviations: NK, natural killer; ZIKV, Zika virus.

### CD14^+^ monocytes prime NK-cell activity during ZIKV infection

Given that peripheral NK cells were activated in ZIKV-infected patients and monocytes are precursors of MDMs, the functional relationship between monocytes and NK cells were assessed. CD14^+^ monocytes were depleted from human primary PBMCs, with an average efficiency of >95% (Supplemental Figure 3). Lipopolysaccharide (LPS; 10ng/ml) was used as a positive control to simulate priming of NK cells by monocytes (18). A significant reduction in the activity of NK cells was observed when CD14-depleted PBMCs were stimulated with LPS compared to LPS stimulation of PBMCs containing CD14^+^ monocytes (Supplemental Figure 4). This effect was evidenced by the reduced levels of the surface markers CD69, CD107a and intracellular IFNγ in depleted cells, verifying that this approach was an efficient strategy for investigating priming of NK cells by CD14^+^ monocytes

PBMCs were then isolated from seven healthy donors and subjected to CD14-depletion before being either infected with ZIKV or stimulated with LPS in parallel to serve as a control to determine activation of NK cells. ZIKV infection in non-depleted PBMCs resulted in high levels of CD107a and IFNγ (Figure 6A) in CD56^+^CD94^+^ NK cells (Supplemental Figure 5) at 36 hpi — an optimal time-point to detect NK-cell priming (19). The opposite effect, however, was observed in ZIKV-infected PBMCs depleted of CD14^+^ monocytes as the levels of both CD107a and IFNγ were significantly reduced (Figure 6B). The levels of CD107a and IFNγ remained high at 72 hpi in nondepleted infected PBMCs compared to depleted infected PBMCs (Supplemental Figure 6). Interestingly, although monocyte depletion did not affect the expression of NK-cell activation receptors NKG2A or NKG2D, a general reduction in NKG2D-expressing NK cells was observed during ZIKV infection (Supplemental Figure 7A and 7B). Surprisingly, the activation marker CD69 was not increased upon ZIKV infection in this study (Supplemental Figure 7C and EV7D). ZIKV viral load was comparable between both conditions (Figure 6C).

**Figure 6:**
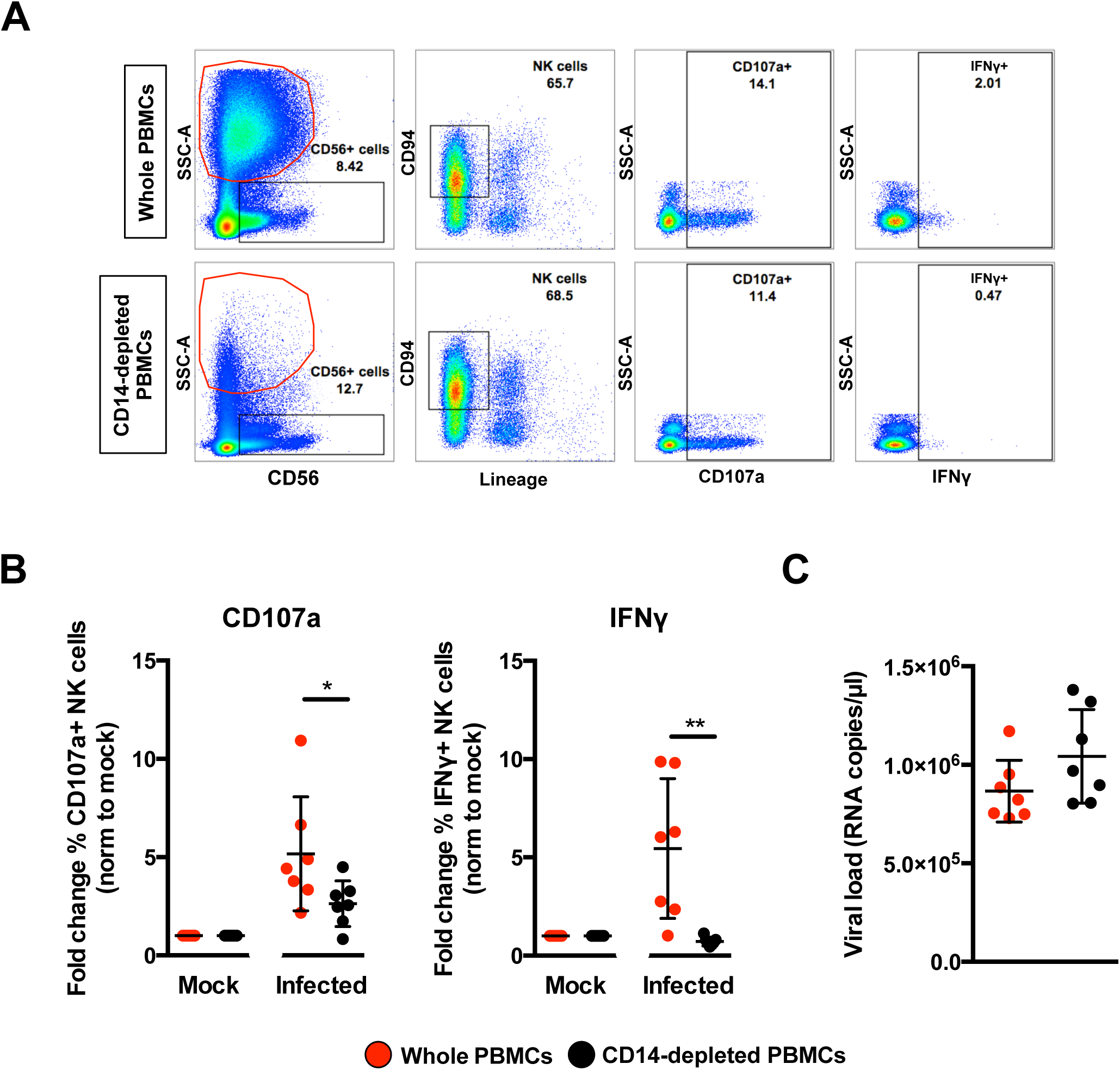
Role of monocytes in NK-cell activity. Full PBMCs and CD14-depleted PBMCs (2×10^6^ cells per infection) were infected with Zika virus (ZIKV) at MOI 10 and harvested at 36 hpi. **(A)** Gating strategy of CD94^+^CD56^+^Lineage-NK cells and their expression of CD69, CD107a and IFNγ. Plots from one representative donor are shown. The red circle indicates the presence or absence of CD14^+^ monocytes. **(B)** Compiled percentages of CD107a and IFNγ-positive NK cells (depicted in **(A))** as normalized to the respective mock sample. **(C)** Viral load in the infected cells. Data shown were derived from seven donors. Lineage markers CD3, CD19, CD20 and CD14 have been included to rule out the presence of non-NK cells. All data are presented as mean ± SD. *\*P* <0.05, *\*\*P* < 0.01, by Mann Whitney *U* test, two tailed. Viral load data was not statistically significant between the two conditions by Mann Whitney *U* test, two tailed. Abbreviations; NK, natural killer; PMBC, peripheral blood mononuclear cell; hpi, hours post-infection.

To delve further into the mechanism, the profile of secreted immune mediators from ZIKV-infected PBMCs was quantified using a 45-plex microbead-based immunoassay (20). Levels of 11 mediators were significantly affected by the depletion of CD14^+^ monocytes (Figure 7A and Supplemental Figure 8A), while 8 mediators were affected upon ZIKV infection (Supplemental Figure 8B). Interestingly, depletion of CD14^+^ monocytes and ZIKV infection did not affect the levels of EGF, IL-9, IL-17A, MIP-1α and MIP-1β (Supplemental Figure 8C). The effect of CD14^+^ monocytes depletion was observed in the levels of SCF and TNFα only after ZIKV infection (Supplemental Figure 8D). Importantly, levels of MCP-1, IL1RA and VEGF-A were affected by both CD14^+^ monocytes depletion and ZIKV infection (Figure 7B). To further investigate the capacity of the cytokine milieus in priming NK cells, freshly isolated human primary PBMCs were then treated with the same culture supernatants from ZIKV-infected PBMCs and CD14^+^ monocytes-depleted PBMCs. Stimulation with culture supernatant from ZIKV-infected non-depleted PBMCs led to slightly more cell death (Supplemental Figure 9A) accompanied by a significant upregulation in expression of CD107a, IFNγ and NKG2D in the CD94^+^CD56^+^ NK cells (Figure 7C and Supplemental Figure 9B), confirming the importance of monocytes in NK-cell priming during ZIKV infection. To rule out priming of NK cells by viruses present in the culture supernatant, a UV-treatment procedure was performed to inactivate the virus, prior to the stimulation assay. Expectedly, while UV-inactivation successfully inactivated ZIKV (Supplemental Figure 10A), it also affected the quality of the cytokines and led to reduced priming of NK cells (Supplemental Figure 10B).

**Figure 7:**
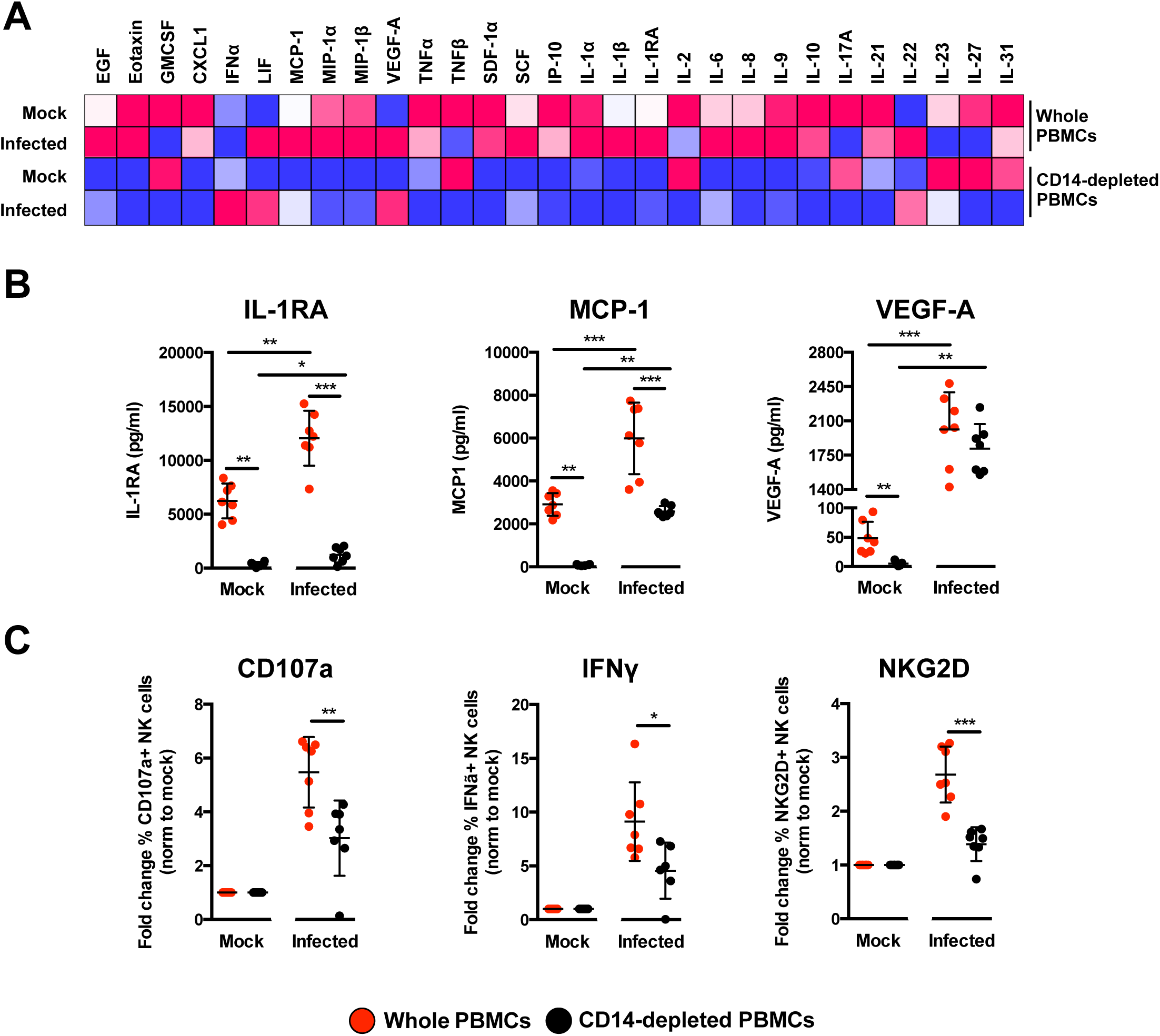
Immune profiling of ZIKV-infected PBMCs. **(A)** Immune mediators in the culture supernatant of ZIKV-infected PBMCs and CD14-depleted PBMCs were quantified with a 45-plex microbeads assay. Concentrations were scaled between 0 and 1. **(B)** Bar-charts of three cytokines, which levels were significantly affected by both the depletion of CD14^+^ monocytes and ZIKV infection. **(C)** Stimulatory capacity of the culture supernatants were further evaluated with freshly isolated PBMCs. Culture supernatant was added in a ratio of 1:10 and cells were harvested at 36 hours post-stimulation. Compiled percentages of CD107a, IFNγ, and NKG2D-positive CD94^+^CD56^+^ NK cells are shown as normalized to the respective mock sample. Data displayed were derived from seven donors. Lineage markers CD3, CD19, CD20 and CD14 have been included to rule out the presence of non-NK cells. All data are presented as mean ± SD. *\*P* <0.05, *\*\*P*<0.01, *\*\*\*p* <0.001, by Mann Whitney *U* test, two tailed. Abbreviations: NK, natural killer; PMBC, peripheral blood mononuclear cell.

## Discussion

Myeloid cells are targets of active ZIKV infection (6-9,21-23) and can elicit immune responses with detrimental outcomes (6,8). Both monocytes and macrophages exhibit extensive heterogeneity (24,25). While it is difficult to obtain tissue-resident macrophages for experimental purposes, human blood is a readily accessible, valuable source of these cells. Transcriptomic profiling of *ex vivo* human blood monocytes and MDMs has revealed marked differences between these cell types (26,27). In this study, human primary monocytes were naturally differentiated into MDMs without any bias for an M1 or M2 macrophage phenotype (28). Given that these cells are targets of ZIKV infection (8), investigations into their cellular immune responses during infection will open avenues to exploit their function for therapeutic benefits.

The level of ZIKV infection (as assessed by the amount of ZIKV antigen and genome copy number) was higher in MDMs than monocytes, which corroborates previous observations (8). Transcriptomic differences between monocytes and MDMs (26,27) would be a plausible explanation for the differential susceptibility of these cells to ZIKV infection. It is also noteworthy that higher ZIKV infection levels were found in purified primary cell populations compared to PBMCs, perhaps due to the presence of other immune subsets in PBMCs that may dampen the overall infection level. ZIKV RNA was detected at the two time-points, 24 and 72 hpi and the virus was present as quasi-species post-infection in human primary myeloid cells. The virus consensus sequence and minor variant mapping revealed an over-representation of transition mutations at highly variable nucleotide positions in the sequence reads. The proportion of these minor variants indicated a shift towards becoming major variants. A recent study that sequenced ZIKV genomes isolated from infected patients provided important information pertaining to ZIKV transmission (29). These data highlighted the degree of divergence in sequenced genomes and placed further emphasis on understanding virus evolution and transmission effectiveness (30). As not all recovered ZIKV RNA samples contained the same mutations, it will be interesting to determine how different host immune responses can lead to ZIKV quasi-species that acquire different combinations of mutations.

ZIKV infection led to the differential abundance of host transcripts mapping to numerous cellular genes in MDMs but not in monocytes, likely due to higher levels of infection observed in MDMs. Furthermore, it has been reported that different donors could account for significant differences in cellular responses (31,32). However, this differential effect does not necessary signify that ZIKV-infected monocytes do not elicit any cellular response to infection, but rather the differences were not measurable by RNA-seq at the read depths used in this analysis. In fact, transcript abundance of numerous genes were different between the mock and ZIKV-infected monocytes, just that the statistical threshold of FDR < 0.05 was not reached and was thus excluded from further analyses. Using IPA data mining, these differentially expressed genes were involved in 133 and 63 canonical cellular pathways (27 of them being shared) in MDMs at 24 and 72 hpi, respectively. The lower number of cellular pathways identified in ZIKV-infected MDMs at the later 72 hpi time-point suggests that certain cellular functions may be shut down post-infection (33). This effect could signify: (1) that the host cells conserve energy to focus only on essential pathways for survival; and/or (2) the host cells have succumbed to ZIKV infection, which leads to transcriptional shutdown in host cells.

Unsurprisingly, the IFN response was the most highly expressed signalling pathway of these common pathways at both time-points because of virus trigger (34). This observation was further complemented by the presence of few other IFN-related pathways. Observations were found for the next two most expressed pathways — pathogenesis of multiple sclerosis and crosstalk between NK cells moDCs cells — both of which involve NK cells. Although ZIKV infection has not been previously associated with multiple sclerosis due to the relatively new disease spectrum, other viral infections such as Epstein-Barr virus (35) and measles virus (36) have been linked.

CXCL9, CXCL10, CXCL11 and CCL5 (identified as the top genes in the pathway) are known chemokines to stimulate NK-cell activation (37,38). The increased transcript abundance of these immune mediators, coupled with others such as IL-15, is a strong indication that ZIKV-infected macrophages are primed to “communicate” with NK cells. Other recent studies have also provided evidence of cross-talk between macrophages and NK cells (18). The increased abundance of TNFSF10 and FAS transcripts in ZIKV-infected MDMs, could indicate priming of NK-cell mediated apoptosis (39). Interestingly, levels of typical NK cell-activating cytokines, such as IL-12 (40,41) and IL-18 (42,43) were not differentially expressed in this study. However, mRNA levels of IL-23 and IL-27, two cytokines belonging to the family of IL-12 (44) with roles in NK-cell activation (45,46) were increased.

Immune-phenotyping of whole blood samples from ZIKV-infected patients revealed the presence of CD69^+^CD56^+^ immune cells (predominantly the CD56^+^ NK cells) (15), suggesting the possible priming of NK cells in ZIKV infection. The involvement of NK cells was thus explored *ex vivo* in human primary PBMCs. Interestingly, *ex vivo* culture alone led to an increase in the basal expression level of CD69 in CD56^+^CD94^+^ NK cells, as previously reported (47). Furthermore, ZIKV infection resulted in reduced levels of CD69, which is a phenomenon also reported for the flavivirus tick-borne encephalitis virus infection in healthy donor NK cells (48). Moreover, NK cells behave differently *ex vivo* and *in vivo* (49), which may explain the different levels of CD69 detected in patients and in *ex vivo* ZIKV-infected NK cells. It was also reported in CD69-deficient mice that the activity of NK cells remains functional (50). High levels of key NK activation markers, including the degranulation marker CD107a and intracellular cytokine IFNγ indicate the higher activation status of NK cells. The activity of NK cells was directly dependent on the presence of CD14^+^ monocytes. ZIKV infection of PBMCs depleted of CD14^+^ monocytes significantly down-regulated the expression of the various NK-cell markers, demonstrating the functional role of monocytes as one of the key players for NK-cell stimulation. The data presented in this study are further supported by a recent publication in which ZIKV patients had high levels of IL-18, TNFα and IFNγ (20) — immune mediators associated with NK-cell function. The usage of SJL mice, which lack NK cells (51), as a model of ZIKV infection also suggested a protective role for these immune cell given that these animals succumbed to cortical malformations (52). Likewise, NK-cell-mediated immune response was significantly increased in healthy volunteers receiving a vaccination for the closely related yellow fever virus (53).

Interestingly, multiplex quantification of secreted immune mediators from *ex vivo* ZIKV-infected PBMCs provided an alternate perspective. IL-18 and IFNγ, two NK-cell related cytokines, were below detection limit. However, freshly isolated PBMCs stimulated with culture supernatants from ZIKV-infected PBMCs resulted in increased priming of NK cells, clearly indicating that the concoction of immune mediators are capable in driving NK-cell activation.

Nonetheless, the, depletion of CD14^+^ monocytes would abrogate this activation as observed by the low levels of MCP-1, IL-1RA, VEGF-A, Eotaxin, GROα, IFNα SDF-1α, IP-10, IL-6, IL-1α, IL-1β, IL-8, IL-21 and IL-10. The reduced levels of MCP-1 could also have a detrimental effect on NK-cell recruitment and priming (37,54), although MCP-1 and VEGF-A have been reported to drive the production of each other (55-57). The high levels of secreted IL1RA from ZIKV-infected PBMCs could also have participated in the increased priming of NK cells, as IL1RA is known to potentiate the effect of IL-2 stimulation of NK cells (58). Thus, the loss of detectable IL-2 after ZIKV infection in CD14-depleted PBMCs would further dwindle NK-cell priming. The presence of other immune mediators such as IL-6, IL-8, IL-10 and IP-10, SDF-1α, GROα, IL-1α and IL-1β in ZIKV-infected non-depleted PBMCs would further provide an inflammatory condition for cellular activation. While the levels of these immune mediators have been reported to be high in ZIKV patients (20), IL-10 and IP-10 have been demonstrated to contribute to cytolysis and activation of NK cells (37,59). Levels of LIF (60), IL-22 (61) and IL-31 (62) were high upon ZIKV infection, indicating their roles in regulating T cells during ZIKV infection (63). T cells can regulate NK-cell activity (64) and monocytes could indirectly mediate NK-cells functions through the T lymphocytes.

To conclude, through a systematic investigative workflow combining approaches exploring host cell transcriptomes and immunoproteomes, it was demonstrated that monocytes and macrophages do not act alone, but in conjunction with other immune cells to orchestrate a series of host immune response and drive disease progression. As such, a comprehensive understanding of immune-cell interaction will have important clinical implications for the design of novel therapeutics that can either dampen down or enhance a response as appropriate.

## Materials and Methods

### Ethics approval and consent to participate

Whole blood samples were collected from ZIKV-infected patients who were referred to the Communicable Disease Centre, Tan Tock Seng Hospital, Singapore. Blood was obtained from patients who provided written informed consent. The study protocol was approved by the SingHealth Centralized Institutional Review Board (CIRB Ref: 2016/2219). Blood samples were collected from healthy donors with written consent in accordance with guidelines from the Health Sciences Authority of Singapore (study approval number: NUS IRB: 10-250).

### Patient whole blood samples

This study utilized whole blood samples obtained from patients admitted to the Communicable Disease Centre at Tan Tock Seng Hospital, Singapore from 27 August to 18 October 2016. Samples included in this study were collected during the acute phase (1-7 pio) of ZIKV infection. These patients were confirmed to be infected with ZIKV by reverse-transcription polymerase chain reaction (RT-PCR) performed on serum and urine samples obtained during their first visit to the clinic. Whole blood samples were collected in EDTA Vacutainer tubes (Becton Dickinson). Whole blood samples were also obtained from healthy volunteers as controls, which were confirmed to be negative for ZIKV RNA by RT-PCR.

### Virus preparation

The ZIKV strain (accession KJ776791) used in this study was originally isolated from the French Polynesia outbreak in 2013 (65). The virus was propagated as previously described (8). Briefly, the virus was propagated by multiple passages in Vero-E6 cells (ATCC; CRL-1587) and pre-cleared by centrifugation before storing at -80°C. The virus titre was determined using standard plaque assays with Vero-E6 cells. Vero-E6 cells were regularly tested for mycoplasma contamination and were grown and passaged in Dulbecco's Modified Eagle Medium (DMEM; HyClone) supplemented with 10% (vol/vol) FBS. UV-inactivation of ZIKV was performed with the CL-1000 UV cross-linker (UVP) at intensity of 100mJ/cm^2^ for 10 minutes.

### Isolation and depletion of monocytes from human PBMCs

Monocytes were prepared from fresh human PBMCs as previously described (8) and by gradient centrifugation using Ficoll-Paque density gradient media (GE Healthcare). Subsequently, monocytes were isolated using an indirect magnetic labelling system (Monocyte Isolation Kit II, Miltenyi Biotec). A direct magnetic labelling system (Human CD14^+^ monocytes isolation kit 2, STEMCELL) was used for depletion of monocytes from PBMCs. The manufacturers’ protocols were strictly adhered to for these procedures.

### Differentiation of monocytes into MDMs

Isolated monocytes were differentiated into MDMs by plating in complete Iscove Modified Dulbecco's Medium (IMDM) (Hyclone) supplemented with 10% (vol/vol) heat-inactivated human serum (HS) (Sigma-Aldrich), which was replaced every 2 days. ZIKV infections were performed on monocytes and MDMs 5 days later, as described below.

#### Virus infection.

ZIKV infections were performed at multiplicity of infection (MOI) 10. Each infection mix consisted of a virus suspension prepared in serum-free IMDM (Hyclone). The cells were incubated with the infection mix at 37°C and allowed to adsorb for 2 h with intermittent shaking before the virus inoculum was removed and replaced with complete IMDM supplemented with 10% (vol/vol) HS (Sigma-Aldrich). Cells were incubated at 37°C until harvesting at 24 and 72 hpi. The harvested cells for downstream total RNA isolation were stored at -80°C. A total of 140 μl of the infected cell suspension was used to quantify the viral load. For assessment of monocyte function in NK-cell activation during ZIKV infection, total human PBMCs and donor-corresponding CD14-depleted PBMCs were infected with ZIKV at MOI 10. In parallel, both PBMC fractions were stimulated with 10ng/ml lipopolysaccharide (LPS; Sigma) as a positive control to measure NK-cell activation. Cells were subsequently treated with 1X Brefeldin (eBioscience) and stained with CD107a (BD Pharmingen) 6 h before harvesting at 36 hpi. The viral load was quantified from 140 μl of the infected cell suspension. Negative controls were cells undergoing the same infection conditions in the absence of infectious ZIKV particles. These controls are referred to as mock-infected samples.

### PBMCs stimulation assay

Fresh PBMCs were isolated as described above and subjected to stimulation with ZIKV-infected culture supernatants in a final ration of 1:10 in fresh IMDM (Hyclone) supplement with 10% (vol/vol) of HS (Sigma-Aldrich). Cells were subsequently treated with 1X Brefeldin (eBioscience) and stained with CD107a (BD Pharmingen) 6 h before harvesting at 36 h for downstream antibodies staining.

### Viral RNA extraction and viral load analysis

Viral RNA was extracted using a QIAamp® Viral RNA Mini Kit (QIAGEN), according to manufacturer's instructions. Quantification of ZIKV NS5 RNA was determined by quantitative real time-PCR (qRT-PCR) TaqMan assay (66) using a QuantiTect® Probe RT-PCR Kit (QIAGEN) in a 12.5 μl reaction volume. All reactions were performed on a 7900HT Fast Real-Time PCR System machine (Applied Biosciences).

### Total RNA extraction

Total RNA was extracted using an RNeasy Mini Kit (QIAGEN) according to the manufacturer's instructions. The extracted total RNA was quantified on a Nanodrop 1000 spectrophotometer (Thermo Fisher Scientific).

### Flow cytometry and antibodies

Detection of ZIKV antigen was carried out in a two-step indirect intracellular labelling process. Briefly, harvested cells were first fixed and permeabilized with FACS lysing solution (BD Biosciences) and FACS permeabilization solution 2 (BD Biosciences), respectively. Antigen staining was then performed with a flavivirus-specific mouse monoclonal antibody (clone 4G2) (Millipore) followed by secondary staining with a goat anti-mouse IgG F(ab’)_2_ antibody (Invitrogen). Cells were then specifically stained for the surface markers CD45 and CD14 (for ZIKV-infected monocytes and MDMs). Dead cells were excluded by staining with the LIVE/DEAD Fixable Aqua Dead Cell Stain Kit (Life Technologies). For PBMCs, surface markers CD45, CD14, CD3, CD19 and CD56 were stained prior to intracellular staining (for ZIKV-infected PBMCs). For patient samples, 100 μl of whole blood was stained for the surface markers, CD45, CD56, CD94, CD16, CD69, CD107a, NKG2D and NKG2A. The stained cells were subsequently incubated with FACS lysing solution (BD Biosciences) to lyse the red blood cells. CD56+ cells were first identified and were subsequently further defined with the CD94 surface marker to give three other subsets - CD56^bright^CD94^hi^, CD56^dim^CD94^hi^ and CD56^dim^CD94^lo^ (16). To specifically assess NK-cell activity *ex vivo,* PBMC fractions were stained for CD107a and various lineage markers (CD3, CD19, CD20 and CD14) (15) in addition to the panel of antibodies used for patient whole blood staining. The usage of lineage markers excludes the presence of non-NK cells in the ensuing analysis. Stained PBMCS were fixed and permeabilized as described above before intracellular staining of ZIKV antigen and IFNγ.

All antibodies used were mouse anti-human and were obtained from BD Pharmingen (CD3, CD19, CD20, CD14, CD69, CD56, CD94, NKG2D, CD107a and IFNγ), Biolegend (CD16 and CD45) and Miltenyi Biotec (NKG2A). Data were acquired on a Fortessa flow cytometer (BD Biosciences) with BD FACSDiva^™^ software. Data analysis was performed using FlowJo version 9.3.2 software (Tree Star, Inc).

### Cytokines quantification using microbead-based immunoassay and data analyses

Cytokine levels in supernatant obtained from mock and ZIKVinfected PBMCs were measured simultaneously using the ProcartaPlex^™^ immunoassay (Thermo Fisher Scientific) detecting for 45 secreted cytokines, chemokines and growth factors including brain derived neurotropic factor (BDNF); Eotaxin/CCL11 ; epidermal growth factor (EGF); fibroblast growth factor 2 (FGF-2); granulocyte macrophage-colony stimulating factor (GM-CSF); growth-related oncogene (GRO) alpha/CXCL1; hepatocyte growth factor (HGF); nerve growth factor (67) beta; leukemia inhibitory factor (10); interferon (IFN) alpha; IFN gamma; interleukin (IL)-1 beta; IL-1 alpha; IL-1RA; IL-2; IL-4; IL-5; IL-6; IL-7; IL-8/CXCL8; IL-9; IL-10; IL-12p70; IL-13; IL-15; IL-17A; IL-18; IL-21; IL-22; IL-23; IL-27; IL-31; interferon-gamma induced protein (IP)-10/CXCL10; monocyte chemoattractant protein (MCP-1/CCL2); macrophage inflammatory protein (MIP)-1 alpha/CCL3; MIP-1 beta/CCL4; regulated on activation, normal T cell expressed and secreted (RANTES)/CCL5; stromal cell-derived factor (SDF)-1 alpha/CXCL12; tumor necrosis factor (TNF) alpha; TNF beta/LTA; Platelets-derived growth factor (PDGF)-BB; placental growth factor (PLGF); stem cell factor (SCF); vascular endothelial growth factor (VEGF)-A; VEGF-D. Preparation of samples, reagents and immunoassay procedures were performed according to manufacturers’ instructions. Data were acquired using Luminex FlexMap 3D® instrument (Millipore) and analyzed using Bio-plex Manager^™^ 6.0 software (Bio-Rad) based on standard curves plotted through a five-parameter logistic curve setting. Levels of BDNF, FGF-2, HGF, NGF, IFN gamma, IL-4, IL-5, IL-7, IL-12p70, IL-13, IL-15, IL-18, RANTES, PDGF-BB, PLGF and VEGF-D were below detection limit and excluded for further analysis. Hierarchical clustering was done using TM4-MeV (http://mev.tm4.org/)

### RNA-seq and differential gene expression analysis

The general approach to RNA-seq and differential expression has been previously described (10,68), and is detailed in brief below.

### RNA-seq

RNA samples were treated with DNase using an Ambion Turbo DNA-free Kit (Ambion), and then purified using Ampure XP beads (Agencourt). The DNase-treated RNA (2 ug) underwent Ribozero treatment using an Epicentre Ribo-Zero Gold Kit (Human/Rat/Mouse) (Epicentre) and re-purified on Ampure XP beads. Successful RNA depletion was verified using a Qubit (Thermo Fisher Scientific) and an Agilent 2100 Bioanalyzer (Agilent) and all of the depleted RNA was used as input material for the ScriptSeq v2 RNA-Seq Library Preparation protocol. RNA was amplified for 14 cycles and the libraries were purified on Ampure XP beads. Each library was quantified using Qubit and the size distribution was assessed using the AATI Fragment Analyser (Advanced Analytical). These final libraries were pooled in equimolar amounts using the Qubit and Fragment Analyser data. The quantity and quality of each pool was assessed by the Fragment Analyser and subsequently by qPCR using the Illumina Library Quantification Kit (KAPA Biosystems) on a Light Cycler LC480II (Roche) according to manufacturer's instructions. The template DNA was denatured according to the protocol described in the Illumina cBot User guide and loaded at 12 pM concentration. Sequencing was carried out on three lanes of an Illumina HiSeq 2500 with version 4 chemistry, generating 2 × 125 bp paired-end reads.

### Bioinformatics Analysis

Briefly, base calling and de-multiplexing of indexed reads was performed using CASAVA version 1.8.2 (Illumina) to produce 30 samples from the five lanes of sequence data in fastq format. The raw fastq files were trimmed to remove the Illumina adapter sequences using Cutadapt version 1.2.1 (69). The option “-O 3” was set so that the 3’ end of any read that matched the adapter sequence by ≥3 bp was removed. The reads were further trimmed to remove low-quality bases using Sickle version 1.200 with a minimum window quality score of 20. After trimming, reads <50 bp were removed. If both reads from a pair passed this filter, each read was included in the R1 (forward reads) or R2 (reverse reads) file. If only one read of a read pair passed this filter, it was included in the R0 (unpaired reads) file. The reference genome used for alignment was the human reference genome assembly GRCh38. The reference sequence was downloaded from the Ensembl ftp site(ftp://ftp.ensembl.org/pub/release77/fasta/homo_sapiens/dna/Homo_sapiensGRCh38.dna_sm.primary_assembly.fa.gz/). The reference annotation was also downloaded from the Ensembl ftp site(ftp://ftp.ensembl.org/pub/release-77/gtf/homo_sapiens/Homo_sapiens.GRCh38.77.gtf.gz/). The annotated file contained 63,152 genes. R1/R2 read pairs were mapped to the reference sequence using TopHat2 version 2.1.0 (70) that employs the mapper Bowtie2 version 2.0.10 (71).

### Differential Gene Expression and Functional Analysis

Mapped reads were further analyzed using EdgeR version 3.3 (72) to calculate normalized counts per million (CPM), identify differentially expressed genes between infected and mock-infected conditions, and compare infected conditions with each other. Correlation and PCA analysis plots were created in RStudio. Heat-maps were generated using GENE-E (Broad Institute;https://software.broadinstitute.org/GENE-E/). IPA was used for gene ontology and pathway analysis. The *P* value associated with each identified canonical pathway was calculated by Fisher's Exact test (right-tailed). The presence of the 27 common canonical pathways was illustrated in a heat-map generated by hierarchical clustering using TM4-MeV (73).

### Identification of ZIKV variants

Bowtie 2 (71) was used to determine the mean sequence coverage. Here, 12 of the 41 samples (including the inoculum) had a mean coverage >10 following alignment with the ZIKV reference genome (accession KJ776791) used in this study. The frequencies of minor variants were calculated using QuasiRecomb (74). Sequences of individual viral proteins were compared to the protein databank using the online NCBI Protein BLAST server(https://blast.ncbi.nlm.nih.gov/Blast.cgi?PAGE=Proteins/).

## Author contributions

FML, DAM, JAH and LFPN designed the study. FML, DL, DAM, JAH and LFPN wrote the manuscript. FML, CTK, CLYP, JJLT, XL, WXY and YXF performed the experiments. FML, DL, XL, YXF, BL, NYR, DAM, JAH and LFPN analysed the data. All other authors were involved in sample collection, processing and analysis, and/or logistical support. All authors read and approved the final manuscript.

## Acknowledgements

The authors would like to thank Siti Naqiah Amrun, Yiu-Wing Kam, Jonathan Cox, Yi-Hao Chan, Guillaume Carissimo, Farhana Abu Bakar, Nicholas Q.R. Kng, Kia-Joo Puan and Nurhashikin Binte Yusof from SIgN for their help in the processing of patient samples. The authors also thank Linda Kay Lee from the Communicable Diseases Centre and Ivy Low, Seri Mustafah and Anis Larbi from the SIgN Flow Cytometry team for their assistance. The authors are grateful to all patients and healthy volunteers for their participation in the study. Finally, the authors would like to thank Laurent Rénia and Insight Editing London for comments and proofreading the manuscript prior to submission. This work was supported by core research grants provided to the Singapore Immunology Network (SIgN) by the Biomedical Research Council (BMRC) and by the BMRC A*STAR-led Zika Virus Consortium Fund [project number: 15/1/82/27/001], Agency for Science, Technology and Research (A*STAR), Singapore. This work was also supported by funds provided by the Medical Research Council (MRC) [project number: MC_PC_15094], and by the HPRU in Emerging and Zoonotic Infections Cross NIHR Strategic Research Fund [project number: ZLKN_11447], United Kingdom. The views expressed are those of the author(s) and not necessarily those of the A*STAR, NHS, the NIHR, the Department of Health or PHE.

